# Distance is not everything in imaging genomics of functional networks: reply to a commentary on *Correlated gene expression supports synchronous activity in brain networks*

**DOI:** 10.1101/132746

**Authors:** Jonas Richiardi, Andre Altmann, Michael Greicius

## Abstract

Our 2015 paper (Richiardi et al., 2015), showed that transcriptional similarity of gene expression level is higher than expected by chance within functional brain networks (defined by functional magnetic resonance imaging), a relationship that is driven by around 140 genes. These results were replicated in vivo in adolescents, where we showed that SNPs of these genes where associated above chance with in-vivo fMRI connectivity, and in the mouse, where mouse orthologs of our genes showed above-chance association with meso-scale axonal connectivity. This paper has received a commentary on biorXiv (Pantazatos and Li, 2016), making several claims about our results and methods, mainly pointing out that Euclidean distance explains our results (“…high within-network SF is entirely attributable to proximity and is unrelated to functional brain networks…”). Here we address these claims and their weaknesses, and show that our original results stand, contrary to the claims made in the commentary.

## 1 Introduction

To a first approximation, all connectivity, like all politics, is local and these two features – nearness and connectivity – are challenging to disentangle. Evidence abounds that the majority of connectivity is local, but this critical attribute of functional anatomy is perhaps most efficiently conveyed in a macaque tracer study by Markov et al. (2012, see in particular Figure 7). Thus, in undertaking our analysis, we were well aware that spatial nearness is correlated with connectivity, which we addressed using a measure of “fine” tissue-tissue similarity derived from an ontological atlas provided by the Allen Institute. The peer-reviewers at Science were also aware of this confound and insisted on an additional level of tissue-tissue similarity correction which our analysis survives (note that Data File S1 of our paper also included information about “coarse” tissue classes (field coarse_tissue_class)).

The commenters suggest that the tissue-tissue similarity correction we applied is inadequate and point out that in the brain samples we used there remains a significant linear correlation between Euclidean distance and correlated gene expression between brain regions. The r-value for this correlation is 0.1. When the coarse tissue-tissue correction is applied, the r-value for the correlation drops to 0.094. In terms of variance explained, this means that slightly less than 1% of the correlated gene expression measure used in our analyses can be explained by Euclidean distance.

We thank the commenters for providing an independent replication of our core results, using the same method and data as in our original paper, and our rebuttal can be summarized in the following 5 points:

1. Our analysis survives correction for Euclidean distance (see details below), applied on top of the tissue-tissue similarity correction we used. Here, we note that there is an intrinsic contradiction in the commenters’ counter-argument that a linear regression of Euclidean distance is inadequate, despite the fact that their critique (see their Figure 1B) is founded on this very same linear correlation.
2. The random clusters generated by the commenters, meant to show the non-specificity of our results, consist of nodes that are roughly twice as close to one another as the nodes in the actual functional networks (see Figure 1 below).
3. We replicated results of our connectivity gene in both a mouse connectivity analysis and a resting-state fMRI connectivity analysis. The commenters did not generate gene lists for any of their random cluster analyses to try and replicate in these or other independent datasets.
4. Euclidean distance correction will wrongly assign “nearness” to two “neurally distant” regions on the crowns of adjacent gyri (see Figure 2). This is essentially the opposite problem of the limitation to tissue-tissue correction that the commenters rightly point out in their figure 1A.
5. Correcting the connectivity of two regions for nearness, using any measure, is bound to dilute the measure of connectivity. That our gene list survives two levels of tissue-tissue similarity correction plus a correction for Euclidean distance and is then replicated in a mouse structural connectivity dataset and a human resting-state fMRI connectivity datasets strikes us as strong support for the conclusion that these genes are important to functional connectivity.

**Figure 1:**
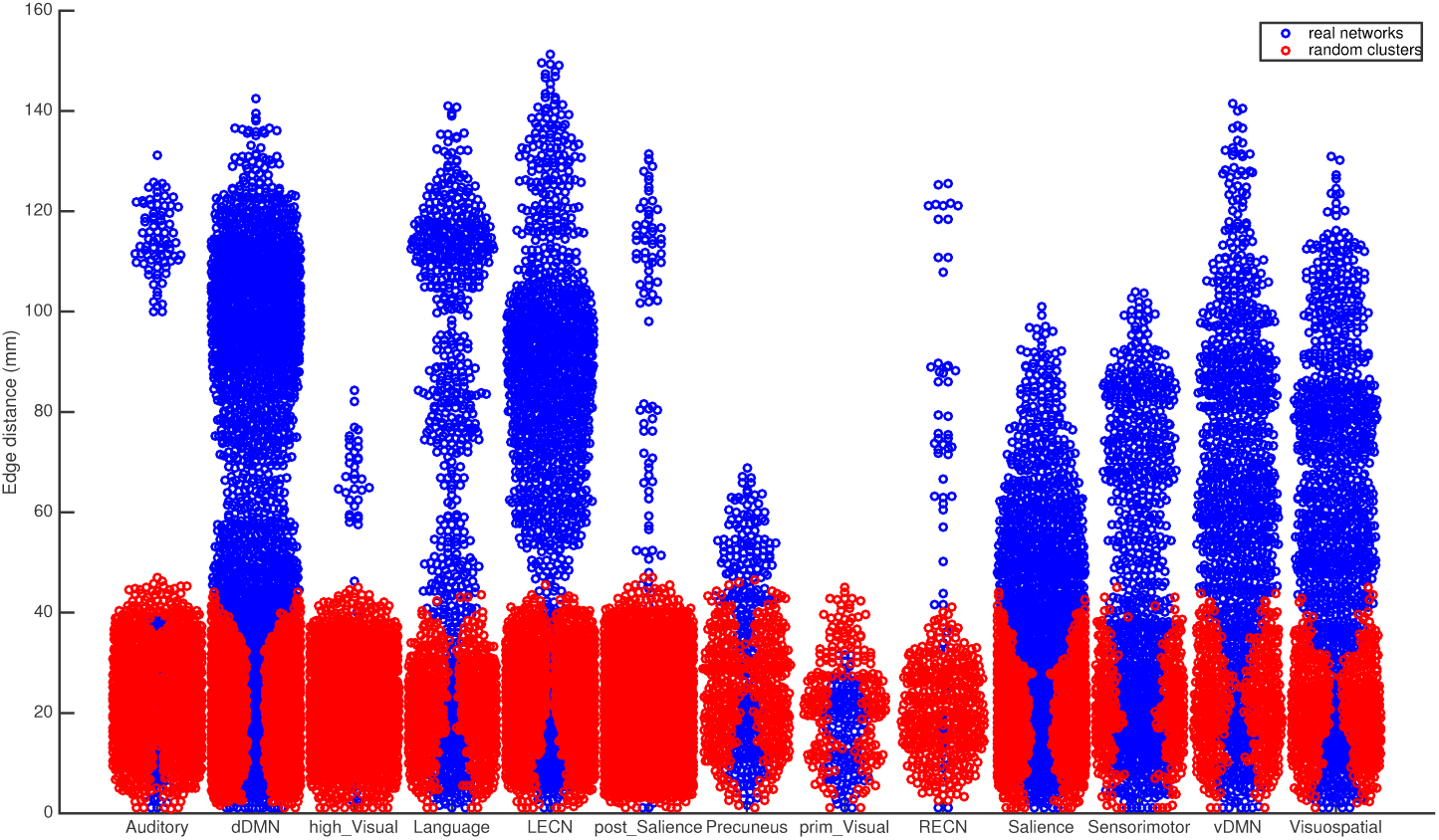
Euclidean distances between samples for the real functional networks (blue) used in our analysis and random clusters (red) generated with the algorithm used by the commenters. The nodes of the random clusters are much nearer to one another than the nodes of the real networks. Random networks of nodes with a similar distribution of Euclidean distances to the real networks would make for a more compelling comparison set but this was not attempted by the commenters.

**Figure 2:**
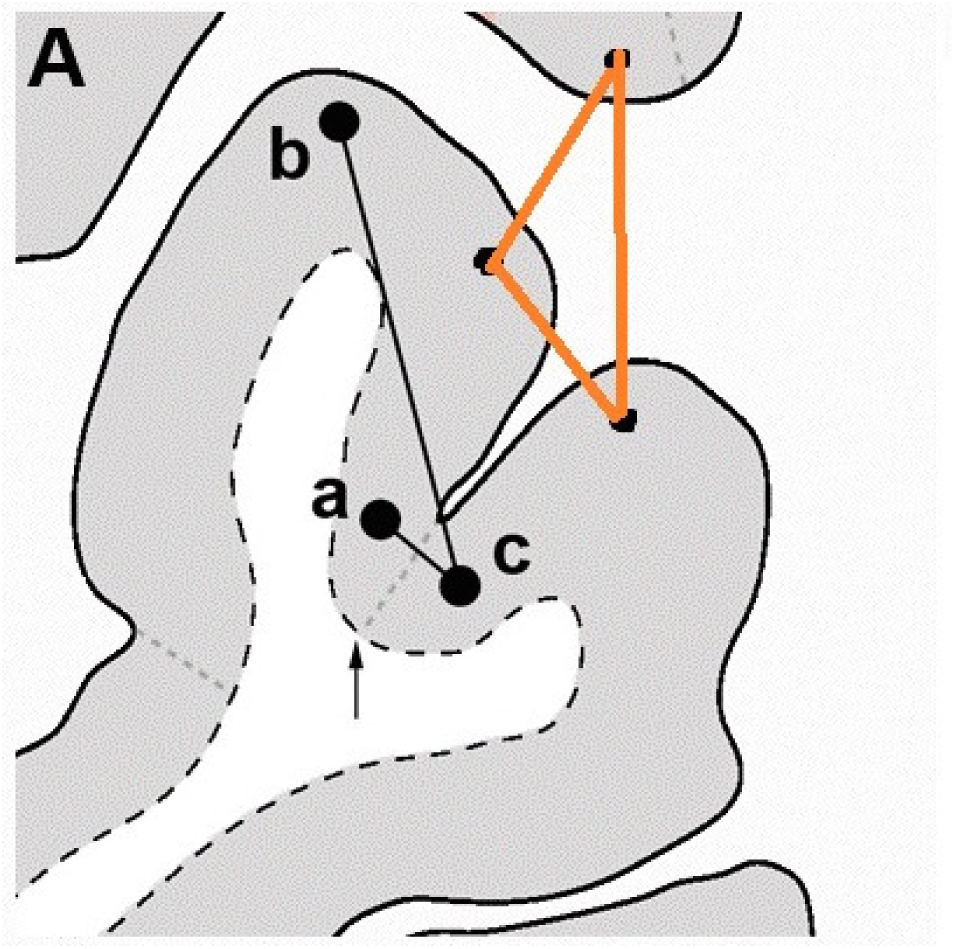
Euclidean distance vs “neural distance”. The orange lines show a case where Euclidean distance would be small even though the tissues may be far neurally. Modified from(Pantazatos and Li, 2016, Figure 1A).

Below, we elaborate on these points.

## 2 Euclidean distance and transcriptional similarity

The commenters say “Even after removing within-tissue edges, there remains an association between tissue tissue correlations and distance (…)”. Indeed such an association remains, although the small effect size (*r*(778123) = *-*0.113*, p <* 0.001) is slightly reduced by removing “fine” tissue-tissue edges (*r*(760785) = *-*0.100*, p <* 0.001), and further slightly reduced by removing “coarse” tissue-tissue edges (*r*(740541) = *-*0.094*, p <* 0.001).

Beyond this well-known association, the main question to address is *whether such an association is sufficient to explain our main results*. We strongly believe it is not, as we show below, because **our results hold both when Euclidean distance is regressed-out**, **and also when a distance-aware permutation testing scheme is used** (Section 4).

To address the Euclidean distance issue, we show that our main result hold when Euclidean distance is regressed out of the transcriptional similarity. This is indeed the case, with different variants on the procedure all yielding significant (at the *α* = 0.05 level) strength fractions. In the first variant, using the vector of transcriptional similarities **s** *∈* ℝ^1577976^, already corrected for intercept, batch IDs, and subject IDs, we can further regress out the vector of corresponding Euclidean distances 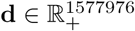 by forming the model **s** *∼* **1** + **d**. We then use the corresponding raw prediction residuals **e** as the *transcriptional similarities with Euclidean distance regressed out*. We then follow our original statistical testing approach, yielding a strength fraction of 0.0053, and a permutation p-value of 0.007. In the second variant, we learn the regression only on the non-negative elements of **s** (leaving 778125 elements) and the corresponding elements of **d**. We then follow the original statistical testing approach, yielding a strength fraction of 0.0061 and a permutation p-value of 0.004. Lastly, in the third variant of the computation we learn the regression only on the non-negative elements of **s** that also are not connecting same tissue classes (leaving 760787 elements), and the corresponding elements of **d**. Here, running the original statistical testing approach yields a strength fraction of SF = 0.0061, and a permutation p-value of 0.001.

However, this approach to linear regression is criticized by Pantazatos and Li, who write “(…) there are two problems with this: 1) The assumption that tissue-tissue correlation strength various linearly with distance is too strong.”

We must point out the contradiction here. Figure 1A in Pantazatos and Li, used to show the dependence between transcriptional similarity and Euclidean distance, also makes use of the linearity assumption since a Pearson correlation result is shown. Pantazatos and Li also write “(…) applying linear regression to adjust for distance (French and Pavlidis, 2011) results in a large negative SF (SF=-0.61, p=1, data not shown)”.

There are several problems with this statement. First, by definition and design, with non-negative edges, SF should never be negative. Second, looking at the code in proc_all.m, lines 289 and following, we see that the following is done:

1. Negative and tissue-tissue edges are masked out (with NaNs) BEFORE regressing out distance. This removes negative-transcriptional-similarity-edges and tissue-tissue edges from the regression, and therefore this only learns the distance-transcriptional similarity relationship on a subset of edges. Since the goal of removing negative edges is to have an non-negative SF, this is not necessary. Running this step is the same as our “third variant” of the Euclidean distance correction procedure above.
2. The original edge mask (negative and tissue-tissue edges) is applied to threshold the resulting matrix. However, the original edges mask does NOT necessarily correspond to negative values in the regressed-out matrix, which is why the resulting matrix can end up with negative edges values and leads to an incorrect computation of SF.

When performed correctly, correcting for Euclidean distance via linear regression shows that our results indeed hold.

We also point out that is has been shown before that genetic effects on brain imaging cortical phenotypes can be sharply different at short ranges: for example Chen et al. (2013, figure 3) show that adjacent brain regions (e.g. superior temporal and posterolateral temporal) can belong to the most separated clusters of genetic influences on surface area.

**Figure 3:**
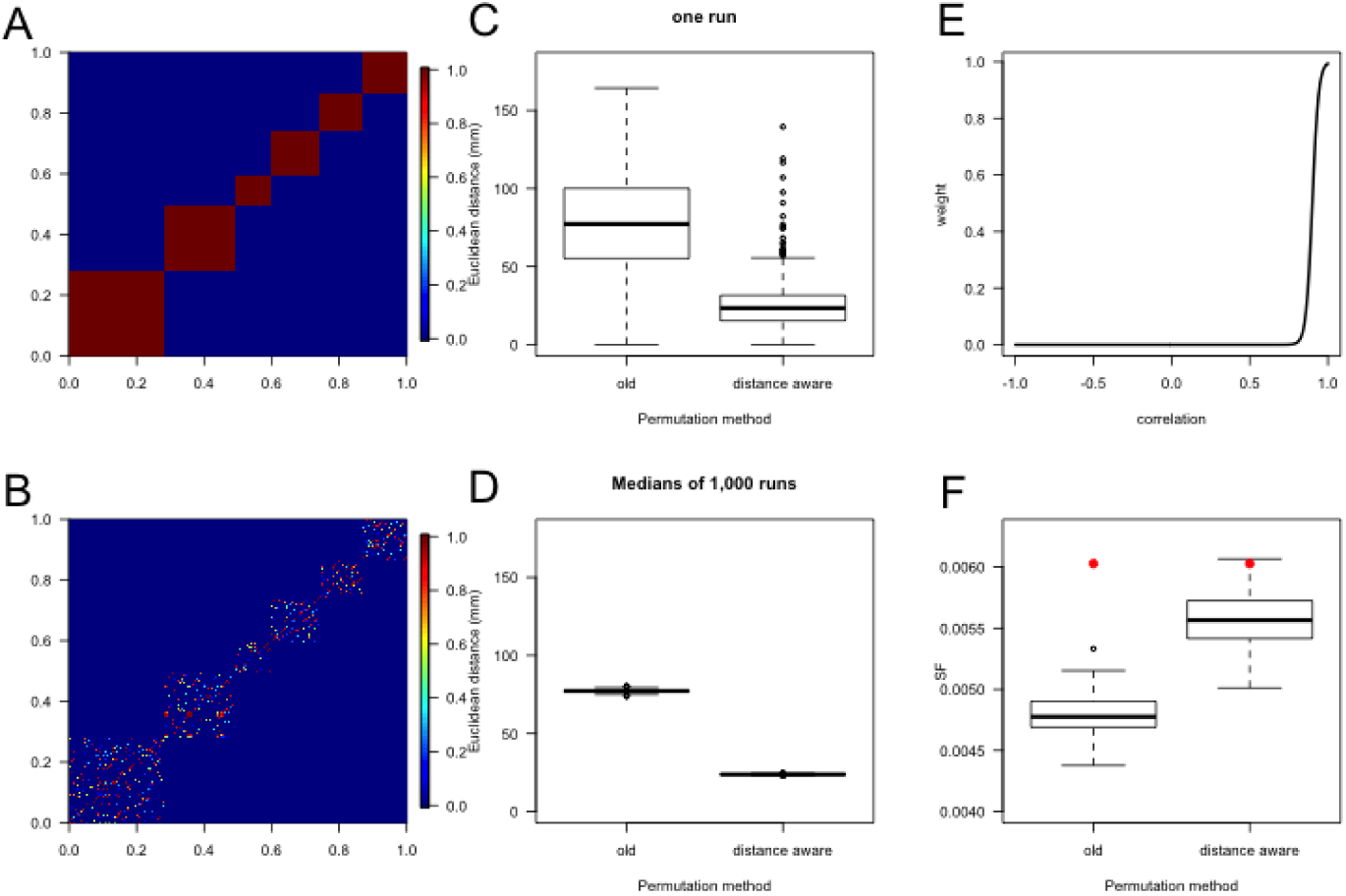
Results with distance-aware permutation procedure. See text for panel details.

## 3 Significance of random clusters

Pantazatos and Li indicate that a significant strength fraction (SF) can be achieved from randomly generated functional networks. We thank the commenters for making this suggestion and probing into the specificity of our result. However, after inspecting the provided code and generating such random functional *networks* it appears that these random networks bear very little similarity to real functional networks: each cluster only comprises regionally close samples. We assume this was a choice to keep the algorithm simple. The downside of this simplification is that real functional networks are more spread out: Figure 1 shows the comparison of within-network edge distances from the real networks and random clusters. This has the effect of overemphasizing the importance and effect of regional similarity, and achieves significance.

## 4 Sample permutation

The commenters raise an interesting point by saying “the null distribution derived in Richiardi et al. is flawed because the permutation strategy assumes all regions are independent and equally exchangeable, which is not true given the spatial autocorrelation and distance bias.”

While this is not literally true (we permute samples within subjects, not across - and this is also implemented correctly in the commenters’ code), we show here that our results hold even when a distance-aware permutation approach is used.

In brief, instead of replacing a sample with an entirely random sample (from the same donor) the distance-aware approach considers the samples’ distance to all other samples during the sampling. Two samples are more likely being swapped if their distances to the remaining samples are similar. In detail, we first compute all pair-wise Euclidean distances between samples. Next, for each pair of samples (within the same brain), we compute the correlation of these distances, hence their similarity is expressed as a value ranging from -1.0 to 1.0. In order to turn this into a weight *w* for the sampling process we perform a sigmoid transformation:

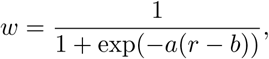

where *r* is the input correlation value. The parameters *a* and *b* control the shape of the curve. Parameter *b* determines which correlation value is mapped to the weight 0.5, the slope *a* is adjusted based on *b* such that the correlation of 1.0 is mapped close to 0.99. For our experiments we selected *a* = 40 and *b* = 0.9. That is, the weighting is biased towards high correlation values: even strong correlations such as *r* = 0.8 receive only a relatively small weight (*w* = 0.0067); correlations of *r* = 0.9 receive *w* = 0.5 (Figure 3, panel E). The weight matrix for the original approach and the distance-aware weight matrix are depicted in Figure 3 panels A and B, respectively. In figure A, the 6 blocks on the diagonal show that permutation is only permitted within each brain. The weight vector is used in R’s sample function to provide a weighted sampling. The approach leads to much shorter distances between selected samples (Figure 3, panel C for one run): the medians for 1,000 permutations are reduced from 75 mm to 25 mm using the distance-aware sampling (Figure 3, panel D). Figure 3, panel F shows the effect of the different sampling procedures on the significance of the strength fraction. With the distance-aware sampling, our results hold with the fine-tissue masking (*P ≤* 0.01). For completeness we repeated this procedure with different values of *a* and *b*: (1) *a* = 20, *b* = 0.8 and (2) *a* = 90, *b* = 0.95. The medians for 1,000 permutations were 32mm and 17mm for (1) and (2), respectively. Likewise, our results held with the fine-tissue masking in both cases, resulting in *P <* 0.01 and *P* = 0.04 for (1) and (2), respectively.

Of note, this distance-aware sampling process is likely over-conservative for our permutation test: due to the distance constraint, many exchanged samples originate from the same region (i.e., a within network sample is replaced with another within-network sample), and thus effectively does not affect the SF.

## 5 Graph thresholding

The commenters say “Applying a cutoff of zero for connections contributing to the SF is not well justified (this applies to the main analyses as well). What, biologically, distinguishes a correlation of 0.1 vs -0.1 other than i.e. noise in the expression vector” First, we must remark on a contradiction in this comment. The whole argument of the commenters is predicated on a correlation value of *r* = 0.1 between transcriptional similarity and Euclidean distance being relevant. So, to return the question to the commenters, what makes *this* value of 0.1 different from zero, other than noise in the expression vector? As we have shown, in this particular case, given the sample sizes, the correlation value is significant, but the effect size is small, and most importantly does not change our conclusions.

Second, we note that any analysis requires some choices. We believe this threshold is a reasonable compromise for two main reasons. Our first reason comes from the literature: our graph representation approach is quite similar to that adopted in “weighted gene co-expression analysis” (Langfelder and Horvath, 2008) for “signed networks”. While we focus on sample-sample interactions, the WGCNA approach focuses on gene-gene networks, but the concerns are similar. In WGCNA, the edge strength of “signed networks” is derived by the “power adjacency function” 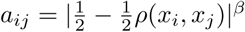, where *ρ*() is the correlation value between two microarray samples, and *β* controls the non-linearity of the transformation. Negative correlations will therefore be squashed down very close to zero with “high” *β* values, an approach which the authors in (Mason et al., 2009) recommend using rather than “unsigned networks”, arguing that diminishing the importance of negative correlations is beneficial. A typical value for *β* (see Langfelder and Horvath, 2008; Mason et al., 2009) for “signed networks” is 12, meaning that negative correlations are essentially transformed to zero edge weight. This handling of negative correlations is equivalent to our approach. The main difference between our approach and the WCGNA approach is in the mapping for correlations between 0 and 1. We use a strictly linear transformation between 0 and 1 (thus acting like a rectifier function), while WCGNA at high *β* values is non-linear and aggressively compresses the high range of correlation values. In our case, since transcriptional similarity is generally high between brain samples in the Allen Institute data, many correlations are high and we preferred not to compress the dynamic range at the high end, hence the choice for a linear mapping for this range of correlation values.

Our second reason has to do with interpretability of our graphs. Our strength fraction statistic is motivated by measures of “communities” in graphs, which essentially quantify how compact and homogeneous subgraphs are. Most of the existing and interpretable measures of modularity in graphs do not deal with negative-valued edges (although some recent work does try and use these too, and modularity optimization algorithms can split the graph into subgraphs with only negative-or positive-valued edges), because it is not obvious that, say, taking the square or absolute value is a good idea – it could lead to the opposite of the desired property of having homogeneous subgraphs.

## 6 IMAGEN validation study

The commenters are claiming that it is unclear whether the gene set enrichment results on the IMAGEN cohort could have been achieved with random draws of gene sets from the background set or the set of “differential stability” genes (see below). As clearly stated in the supplement of our published article (section “IMAGEN validation procedure”) we did test whether gene sets of size 136 sampled from all genes or the background set only achieve similar enrichment scores. For each of the two scenarios we conducted a permutation test, drawing 10,000 sets of genes of size 136, and compared the resulting enrichment scores to the one of our real list. In both cases, the real list achieved a significantly better enrichment score (*P <* 4 *×* 10^*-*4^ for all genes and and *P* = 0.0059 for background only).

In terms of the DS genes of Hawrylycz et al. (2015), we first note that this was not available at the time of writing, having been published online on the 16th of November, 2015. Then, in the supplementary discussion to their comment, the commenters write that “… *>*75% of these 136 consensus genes are in the top 10% of genes found to have consistently high region-to-region variability (so called differentially stable, DS genes) across the cortex…”, but this is to be contrasted with what is reported in Hawrylycz et al. (2015, page 11), where the authors state “…[the 136 genes] constitute only a minority (15%) of the highest cortical 5th percentile DS genes.”. Further, they also mention “… higher cortical DS of a gene predicts a stronger correlation between functional connectivity and cortical gene expression pattern …; the correlation with brain-wide DS alone was much weaker …, indicating that the conserved cortical genes correlate most strongly with cortical functional connectivity”.

## 7 Mouse validation study

In the supplementary discussion, the commenters say “the results from mouse tractography data … does not make any adjustment for spatial proximity, and is likely also confounded by spatial proximity.”. We show here that this is not the case. We re-ran our permutation procedure, but substituting for each transcriptional similarity matrix (both original and those computed from a random permutation of 57 genes) a distance-regressed version of transcriptional similarity matrix, obtained from the regression residuals of the linear model **s** *∼* **1** + **d**, where **s** is transcriptional similarity and **d** is Euclidean distance. We then follow the same test procedure as in the original paper.

The results, shown in Figure 4, show that the results survive correction for Euclidean distance, with a significant p-value.

**Figure 4:**
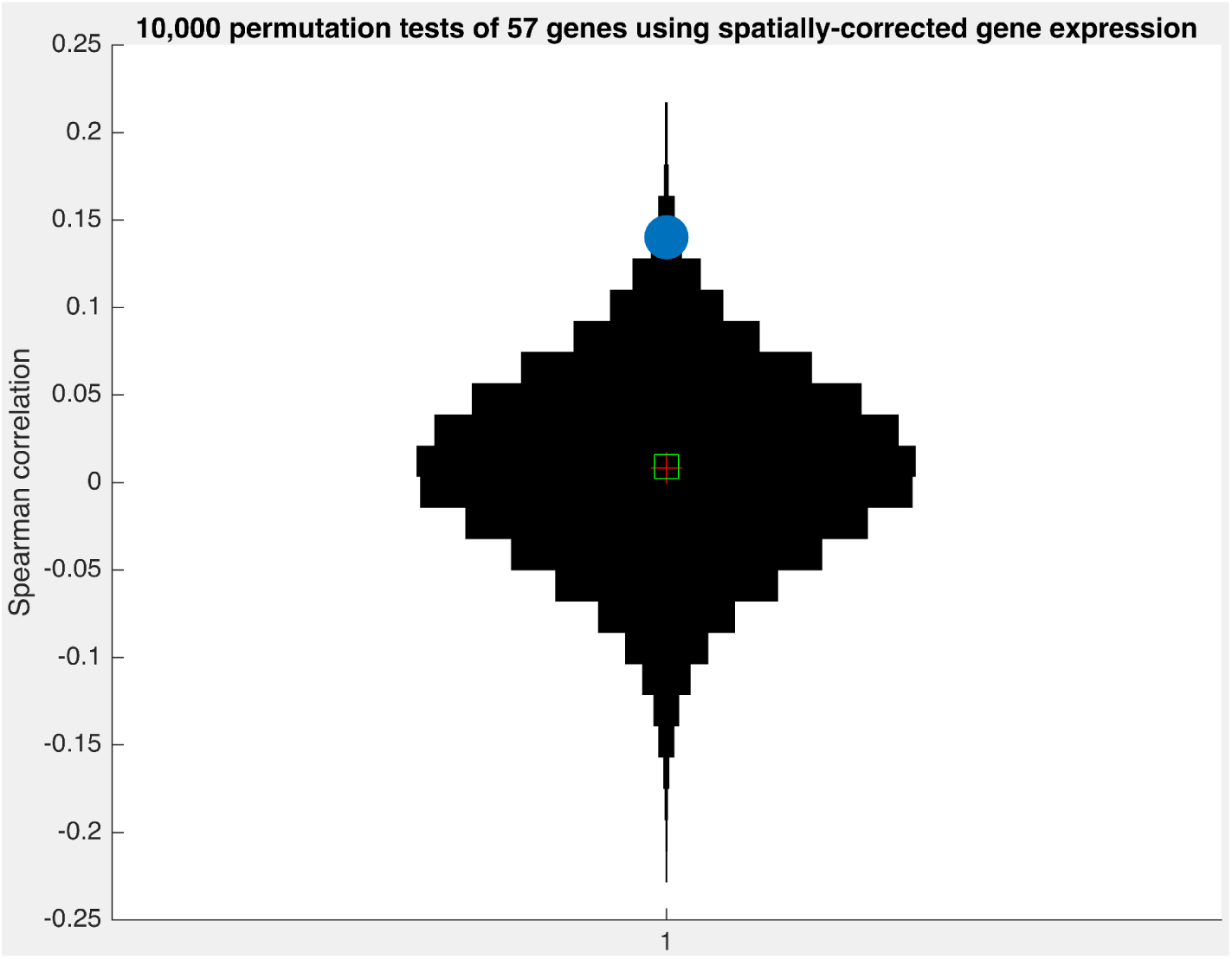
Mantel test statistic distribution for 10,000 permutations of 57 genes (black violin plot) and original set of genes (blue dot) using distance-corrected gene expression data. The original set of genes is significantly more associated with axonal meso-scale connectivity in the mouse than chance (P = 0.009).

## 8 Independent support for our results

Lastly, we note that several published studies have found results consistent with our own.

Recently, Krienen et al. (2016) have shown that regions with similar transcriptional profiles across macro-scale functional networks may be more likely to be connected. In this study, our gene set is explicitly used in (and called the “human/-mouse connectivity set”). Among other issues, this paper discusses distance effects on transcriptional similarity at length, and clearly mentions that this relationship depends on functional networks: “…spatial proximity does not uniformly affect the tendencies of brain samples to have similar or dissimilar transcriptional profiles: at equivalent spatial distances, paralimbic and certain associational networks are more likely to have similar expression profiles to each other than to auditory, visual, or somato/motor profiles.” This is also the case for our gene set.

Using a two-functional-networks model of resting-state activity (visual-sensorimotor-auditory and parieto-temporo-frontal), Cioli et al. (2014) have shown that spatially overlapping differences in gene expression (including ion channels and neurotransmitter release genes) can reliabily predict network membership.

Using RNA-Seq of around 50 human brains in 10 neocortical regions, Wang et al. (2015) showed that fractional amplitude of low-frequency fluctuations (fALFF) in the default-mode network (one of our functional networks) is more correlated with the expression of 38 genes than expected by chance. Further, these authors found that 4 of our genes overlap with their list, explaining more than 10% of the variation in fALFF.

Also recently, Fulcher and Fornito (2016, Figure S3) have shown that bidirectionally connected regions of the mouse brain are more transcriptionally similar than unidirectionally connected regions, and that both are more transcriptionally similar than unconnected regions. This pattern is significantly conserved both with and without exponential distance correction. Because functional connectivity overlaps importantly with structural connectivity, this is another indication in our direction. More generally, there are a number of studies supporting non-trivial links between micro- and macro-scale features of brain organization (Heuvel and Yeo, 2017), and recent work has shown that at the micro level, spatial proximity is not sufficient to explain axonal connections, cell identity being a better guide (Kasthuri et al., 2015).

## 9 Conclusion

Having performed a number of additional analyses in this response, we stand by our original results.

## Acknowledgements

We thank Mario Chavez for his useful comments on this response. All remaining mistakes are our own.

## Source code

Source code related to these and original experiments is being released at: https://github.com/jonasRichiardi/imaginggenomics.

